# CDK4/6 inhibitors promote PARP1 degradation and act synergistically with PARP inhibitors in non-small cell lung cancer

**DOI:** 10.1101/2024.07.07.602389

**Authors:** Carlos M Roggero, Anwesha B Ghosh, Anvita Devineni, Shihong Ma, Eliot Blatt, Ganesh V. Raj, Yi Yin

**Affiliations:** Department of Urology, c, Dallas, TX 75390, USA; Department of Pharmacology, University of Texas Southwestern Medical Center, Dallas, TX 75390, USA

**Keywords:** DNA damage repair, CDK4/6 inhibition, PARP inhibition, patient-derived explant, non-small cell lung cancer

## Abstract

Despite the widespread deregulation of CDK4/6 activity in non-small cell lung cancer (NSCLC), the clinical trials with CDK4/6 inhibitors (CDK4/6is) as a monotherapy have shown poor antitumor activity. However, our preclinical studies have revealed a significant potential for CDK4/6is to collaborate by influencing DNA damage repair pathways during radiotherapy. Given the considerable upregulation of PARP1 expression in NSCLC, we analyzed the efficacy of combined PARP and CDK4/6 inhibition in NSCLC models. Our findings demonstrate that CDK4/6is synergize with PARP inhibitors (PARPis) to inhibit the clonogenic growth of RB-proficient NSCLC models. This synergy is associated with increased accumulation of DNA damage, interrupted cell-cycle checkpoints, and enhanced apoptotic cell death. We showed that CDK4/6is mechanically promote PARP1 protein degradation, leading to decreased availability of DNA repair factors involved in homologous recombination and suppression of DNA repair competency. Furthermore, we showed that PARP trapping is required for this synergy. We then confirmed that combining PARPi and CDK4/6i blocked the growth of NSCLC xenografts in vivo and patient-derived explant models ex vivo. These findings reveal a previously uncharacterized impact of CDK4/6i on PARP1 levels in RB-proficient NSCLC models and the requirement of PARP trapping to render synergy between CDK4/6i and PARPi. Our research suggests that combining CDK4/6i with PARPi could be a promising therapeutic strategy for patients with RB-proficient NSCLC, potentially opening up new and more effective avenues for treatment.

## 1. Introduction

Aberrant activity of cell cycle regulator proteins is one of the hallmark somatic events in non-small cell lung cancer (NSCLC) (1, 2). In normal cells, the transition from G1 to S phase during cell cycle progression is tightly regulated by phosphorylation of the retinoblastoma protein tumor suppressor (RB) through the cyclin-dependent kinase (CDK) 4/6-D-type cyclin (Cyclin D) complex. CDK4/6 interacts with cyclin D, forming functional complexes that phosphorylate RB. Meanwhile, p16INK4A (p16) encoded by the CDKN2A gene inhibits the functioning of the CDK4/6-Cyclin D complex by directly binding to CDK4 and CDK6, thereby preventing phosphorylation of RB and inducing cell cycle arrest (3). In NSCLC patients, p16 is frequently mutated, deleted, or epigenetically silenced (4, 5), resulting in an enhanced phosphorylation of RB and uncontrolled cell cycle progression (1, 2, 5). Because CDK4/6 activity is typically deregulated and overactive in NSCLC (4–6), targeting CDK4/6 is considered an attractive therapeutic strategy against NSCLC. While CDK4/6 inhibitors (CDK4/6i; palbociclib and abemaciclib) have shown some efficacy in preclinical models of NSCLC, results from clinical trials using CDK4/6i as single-agent treatment have been disappointing. A sub-study (S1400C) from the Lung-MAP (SWOG S1400) clinical trial showed partial responses with palbociclib monotherapy in only 2/32 genomically stratified patients and did not meet the criteria for advancement to phase III testing (7). The failure of CDK4/6 monotherapy trials has prompted studies on the effect of the combination of CDK4/6i with other therapies (8, 9).

NSCLC has remarkably high genomic instability, with an exceptionally high number of genomic aberrations, indicating DNA repair deficiency (5). Although DNA damage and genomic instability are possible contributory factors to lung cancer pathogenesis and the development of resistance to different therapies, they also present vulnerabilities that can be exploited therapeutically. The DNA double-strand breaks (DSBs) are one of the most cytotoxic forms of DNA damage. DSBs can be induced by ionizing radiation (IR), chemotherapeutic drugs, oxidative stress, and replication fork collapse (10). DSBs can be repaired by two main pathways: the classical non-homologous end joining (NHEJ) and homologous recombination (HR). One of the first signaling events upon DNA damage is the recruitment of poly (ADP-ribose) polymerase-1 (PARP1) to sites of DNA damage, particularly DNA strand breaks. PARP1 is crucial to multiple DNA repair pathways, including DNA base excision (BER), single strand (SSB), and DSB repair. Notably, among the PARP isoforms endowed with the capacity of poly ADP-ribosylation, the mRNA level of PARP1 is the only one that significantly upregulated in adenocarcinomas and squamous carcinomas of the lungs as compared with normal adjacent tissues (11). In addition, in patients with NSCLC, a high abundance of poly (ADP-ribose) (PAR) indicates poor prognosis (12), suggesting that targeting PARP1 activity is clinically relevant for NSCLC. PARP inhibitors (PARPis) are the first drugs to exploit the concept of synthetic lethality for DNA damage repair (DDR) defects in the clinic. PARPis have shown remarkable activity in HR-deficient preclinical models and attractive response rates in BRCA1 or BRCA2 mutation carriers in different tumor types. Even though HR defects are enriched in tumors that respond to PARPi, not all PARPi responders have readily apparent DDR aberrations, nor do all tumors harboring mutations/deletions in DDR genes respond to PARPi. Importantly, clinical studies assessing PARPi efficacy either in combination with chemotherapy or as maintenance treatment have failed to yield any significant benefit in NSCLC tumors with or without BRCA mutations (13, 14), which has significantly limited the clinical utility of PARPi in NSCLC.

The cyclin D-CDK4/6-RB pathway is also involved in DDR (15), which makes CDK4/6i a perfect candidate for tumor radiosensitization. Significant preclinical data have demonstrated the radiosensitization effects of CDK4/6is via inhibiting DDR, enhancing apoptosis, and blocking cell cycle progression. A preclinical study showed that abemaciclib and IR had a good radiosensitization effect on tumor cells in proliferative and plateau-phase and tumor xenografts but had little radiosensitization effect on normal cells and improved the radiation sensitivity of NSCLC *in vitro* and *in vivo* (16). Abemaciclib inhibited IR-induced DNA damage repair and caused RB-dependent cell cycle arrest.

In this study, we investigated the efficacy of CDK4/6is (palbociclib or abemaciclib) and PARPis (olaparib or rucaparib) combination therapy in NSCLC models and explored the mechanistic underpinnings of our findings.

## 2. Materials and Methods

### 2.1. Cell culture and drug treatments

Human NSCLC cell lines A549, H460, H1437, and H2009 were provided by Dr. John D. Minna (The University of Texas Southwestern Medical Center, Dallas, TX). Human prostate cancer cell line DU145 was obtained from the American Type Culture Collection (ATCC). All cell lines were cultured in RPMI1640 medium supplemented with 10% fetal bovine serum (FBS) (HyClone, Hudson, NH, USA), 10☐mM HEPES, 1☐mM sodium bicarbonate, 100 U/ml penicillin, and 100 μg/ml streptomycin in a humidified incubator at 37☐°C with 5% CO2. Cell lines were obtained during 2013-22, and cultures passed for <3 months after thawing a given frozen vial. The passage range of 5-20 was used in our experiments. Palbociclib, abemaciclib, olaparib, rucaparib and veliparib were purchased from Selleck Chemicals (Houston, Texas, USA). PARP1 degrader rucaparib-AP6 was provided by Dr. Yonghao Yu (The University of Texas Southwestern Medical Center, Dallas, TX) (17). Drug stock solutions were made in dimethyl sulfoxide (DMSO) at 10 mmol/L or 100 mmol/L. The stock solutions were stored at-80°C in the dark and diluted in culture medium immediately before use.

### 2.2. Irradiation

Cells were irradiated with γ-rays generated by a Mark 1 137Cs irradiator (J.L. Shepherd and Associates) at the doses denoted in the figures.

### 2.3. Ex vivo cultures of tumor explants

For patient-derived explants (PDE), tumor tissues were obtained from patients undergoing surgical resection from the University of Texas Southwestern Medical Center and Partner Institutions. Tumor samples were processed for ex vivo cultures, adapting a previously published protocol (18). Under this protocol, tumor tissues not needed for pathological diagnosis were placed in chilled PBS typically within 30 min of surgical resection and arrived in the laboratory 30-60 min later. Specimens were dissected into approximately 1-2 mm^3^ pieces and cultured in quadruplicate, as described previously (18). PDEs were cultured for 72 h in the presence or absence of vehicle control (DMSO), palbociclib (1 μM), or olaparib (2 μM). Following treatments, tumor fragments were then formalin-fixed and paraffin-embedded. For xenograft-derived explants (XDEs), athymic nude mice (nu/nu, 5-6 weeks old) were injected (1× 10^6^ cells in 100 μL 50% Matrigel) subcutaneously (s.c.) into the right posterior flanks. Mice were sacrificed when tumor size reached 500 mm^3^. After excising the xenografts aseptically from mice, tumors were cut into 1-2 mm^3^ tissue fragments and similarly cultured as XDEs to PDEs.

### 2.4. Clonogenic survival assays

Colony formation assay was performed by seeding 500-1,000 cells in six-well plates (depending on the growth rate of each cell line). The culture medium was replaced the next day with a medium containing the appropriate drug(s), and cells were incubated for 10-14 days. Then, the culture medium was removed, and cells were washed in PBS, fixed in 10% formalin, rewashed in PBS, and stained with 0.1% crystal violet aqueous stain. Cells were washed until all stain was removed, air-dried, and scanned. Colonies were quantified using ImageJ/Fiji software (https://fiji.sc/). We quantified the % area stained within each well in Image J software. Adjustments were made to the regions of interest for artifacts, and the quantification results were finalized. Due to the variability in the number of colonies in the control group in biological replicates, we could not combine quantification from separate experiments.

### 2.5. Cell viability assays

Cells were seeded into 96-well plates (500-2,000 cells per well). 24 h after seeding, serial CDK4/6i or PARPi dilutions were added to cells to final drug concentrations ranging from 0 to 10 µM. Cells were then incubated for 4-5 days, and cell viability was measured using Hoechst staining for DNA. Briefly, media from cells plated in 96 well formats was aspirated, and cells were washed once with PBS, followed by adding 100 µL dH20/well, and then stored at-20 degrees. The next day, plates were thawed at room temperature, followed by the addition of 200 µL of freshly diluted Hoecsht stain (1:1000 in TNE buffer containing 1mM EDTA, 2M NaCl, 10mM Tris at pH 7.5). Relative survival in the presence of different inhibitors was normalized to the untreated controls after background subtraction.

### 2.6. Drug synergy and combination index

All drug concentrations were simultaneously combined in a non-constant ratio, or constant ratio (1:1), and the combination index (CI) values were calculated by the Chou-Talalay method with CompuSyn software (19, 20). In general, combinations with a CI value < 1.0 are considered synergistic.

### 2.7. Flow cytometry

Cell-cycle analysis was performed using propidium iodide (PI) flow cytometry. Cells were cultured at 15,000 cells/cm^2^ and treated with vehicle control (DMSO), palbociclib (1µM), and olaparib (2.5 µM) alone or in combination for the indicated times. Cells were trypsinized and washed with 3% FBS in PBS, then fixed with 75% ethanol in PBS. Cells were then collected by centrifugation and washed with 3% FBS in PBS followed by PI (Millipore Sigma, catalog P4170) staining (25 μg/mL PI, 500 μg/mL RNAse, 3.6 mM sodium citrate) overnight at 4°C. Cell cells were resuspended in 300 μL fresh 25 μg/mL PI the following day. The cell cycle phases were assessed using the BD FACS LSRFortessa (BD Bioscience, New Jersey, USA) and FlowJo version 10.0. The percentage of apoptotic cells was measured using the Annexin-V/PI double-staining apoptosis assay. Cells were cultured at 1,500 cells/cm^2^ and treated with vehicle control (DMSO), palbociclib (1 μM), and olaparib (2 μM) alone or in combination for the indicated times. According to the manufacturer’s protocol, 100,000 cells per condition were collected and stained with Alexa Fluor 647 Annexin V (BioLegend, catalog 640912). The number of apoptotic cells was measured using a FACS LSRFortessa flow cytometer, and the percentage of early (Annexin-V positive/PI negative) and late (Annexin-V/PI positive) apoptotic cells was calculated using FlowJo version 10.0.

### 2.8. Quantitative immunofluorescence of DNA damage associated foci

Cells grown on coverslips were fixed using 4% Paraformaldehyde solution in PBS and permeabilized using 0.2% Triton X-100 in PBS for 20☐ min. Blocking was achieved using 2.5% horse and goat serum in PBS. Primary antibodies [anti-p-ATM (Abcam ab36801) 1/300, anti-γH2AX (Millipore 3292608) 1/300, anti-53BP1 (Novus Biologicals NB100-304) 1/300, anti-p-DNA-PKcs (Abcam ab18192) 1/300, anti-RAD51 (Abcam ab63808), anti-BRCA1(Santa Cruz sc-6954)] were diluted in blocking buffer and incubated overnight at 4 ☐°C. Secondary antibodies conjugated to fluorescein or Texas Red (1/500) were used to visualize the primary antibodies. Images were acquired with a camera mounted on a Nikon Eclipse E600 fluorescence microscope or Lionheart™ FX Automated Microscope. To quantify DNA damage associated with immunofluorescent foci, cells with 5 or more foci (to reflect an estimate of induced damage, based on most cells at 0☐Gy having 0-4 foci) were scored as positive. The number of cells containing foci (>4) was determined by manually counting 150-200 cells across 5 or more high-power (200×) microscope fields. Percentages of foci–positive cells were calculated.

### 2.9. Tumor tissue immunohistochemistry staining

Tumor tissue immunohistochemistry staining has been described previously (18). Tissue fragments were fixed overnight in 10% formalin and placed in embedded cassettes. After dehydration in 70% ethanol, formalin-fixed tumors were processed using automated standard procedures and subsequently embedded in paraffin. Four-micrometer tissue sections obtained with Leica microtome were mounted on coated microscope slides. For immunofluorescent staining, sections were deparaffinized using xylene and hydrated with declining ethanol concentrations. Target antigen retrieval was performed using Vector Antigen Retrieval buffer (pH 6.0), heated to 100 °C for 20 min. Cells were permeabilized using PBS with 0.2% Triton X-100 for 20 min. Blocking was achieved using PBS with 2.5% horse and goat serum. Sections were incubated with the appropriate primary and secondary antibodies, developed using 3 3′ diaminobenzidine chromogen and counterstained with hematoxylin. Images were acquired with a camera mounted on a Nikon Eclipse E600 fluorescence microscope or Lionheart™ FX Automated Microscope.

### 2.10. Western blotting

Immunoblotting was performed as previously described (21). Briefly, whole cell samples were lysed in NP40 extraction buffer containing 50 mmol/L Tris-Cl, pH 7.6, 150 mmol/L NaCl, 5 mmol/L EDTA, 1% NP40, 2 mmol/L dithiothreitol (DTT), 1× Phosphatase Inhibitor Cocktail (Sigma), and 1× Protease Inhibitor Cocktail (Roche). 10-50 μg total protein was fractionated by SDS-PAGE and transferred to PVDF or nitrocellulose membranes (Millipore). Blocking was achieved with 5% milk in Tris-buffered saline with 0.1% Tween-20 (TBST) at room temperature for 1h, and blots were then incubated in primary antibodies [anti-PARP1 (Cell Signaling 9542, 1/750), anti-PAR (Enzo Life Sciences, LX-804-220-R100, 1/500), anti-γH2AX (Millipore 3292608, 1/1000), anti-β-actin (Cell Signaling, 1/3000)] diluted in 3% BSA in TBST overnight at 4°C. Following three washes with TBST, blots were incubated with horseradish peroxidase-conjugated goat anti-rabbit IgG (H + L) and goat anti-mouse (H + L) (Jackson ImmunoResearch) as secondary antibodies (1/1000) for 2h at room temperature and then washed thrice with TBST. Pierce™ ECL (Thermal Scientific) or ECL™ prime (GE Healthcare) were used to visualize the substrate using the Bio-Rad ChemiDoc MP imaging system.

### 2.11. Quantitative reverse transcription-PCR

Total RNA was isolated using the RNeasy mini kit according to the manufacturer’s instructions (Qiagen, Valencia, CA) and reverse-transcribed using the cDNA synthesis kit (Bio-Rad, Hercules, CA). Quantitative PCR was performed using iQ SYBR Green Supermix (Bio-Rad) on an iCycler thermal cycler (Bio-Rad) with a denaturation step at 95°C for 3m followed by 40 cycles of amplification at 95°C for 30s, 60°C for 30s, and 72°C for 60s. The following primers were used for PARP1 (forward, TGGAACATCAAGGACGAGCT; reverse, CATCGCTCTTGAAGACCAGC). PARP1 expression was normalized to 18S rRNA (forward, TGTGCCGCTAGAGGTGAAATT; reverse, TGGCAAATGCTTTCGCTTT). Fold changes in mRNA expression levels were calculated using the comparative CT method.

### 2.12. Comet assay

The comet assay was performed as previously described (21). Cells were subjected to different treatments for the indicated times, harvested, and re-suspended in ice-cold PBS (without Mg^2+^ and Ca^2+^). The amount of damaged DNA was visualized by single-cell agarose gel electrophoresis under alkaline conditions following the manufacturer’s protocol. 100 µl/well of diluted Vista Green DNA dye was applied onto slides. Comet images were taken by Lionheart LX Automated Microscope (Agilent BioTek) using a FITC filter and quantified using OpenComet (22). The level of DNA damage was presented as a percentage of DNA in the tail.

### 2.13. PARP Trapping assays

To prepare nuclear-soluble and chromatin-bound subcellular fractions, semiconfluent human A549 cells with 10 mL medium in a 10 cm dish were treated with indicated drugs for 4 h, and then cells were collected. Chromatin fractions were prepared using Thermo Scientific subcellular protein fractionation kits (cat. no. 78840) per the manufacturer’s protocol. Samples were normalized for protein concentration by micro-BCA assay (Pierce BCA Assay Kit, Thermoscientific) and analyzed by immunoblotting (anti-PARP1 cat. no. 9542 and anti-H3, cat. no. 3638; Cell Signaling).

### 2.14. Tumor growth

Male athymic nude mice (nu/nu, 5-6 weeks old) were injected s.c. (1×10^6^ cells in 100 μL 50% Matrigel) into the right posterior flanks. Tumor volume was determined using a caliper, measuring the shorter and larger diameters. Assuming the tumor shape to be ellipsoidal, we applied the formula (short diameter × short diameter × large diameter)/2 to estimate the volume. When subcutaneous tumors reached 100-150 mm^3^ in volume, mice were randomized to different treatment groups, including at least 4-5 tumors per group. Mice were treated with palbociclib alone, olaparib alone, or two drugs in combination for the indicated time when tumor size reached 200 mm^3^. Palbociclib was dissolved in 50 mM sodium lactate solution and administered via oral gavage at 100 mg/kg/day. Olaparib was dissolved in 10% hydroxypropyl-β-cyclodextrin for intraperitoneal administration and dosed at 50 mg/kg/day. Mice were euthanized when tumor volumes reached a maximum of 2,500 mm^3^ or when skin ulcerations occurred.

### 2.15. Study approval

De-identified tumor tissues were obtained from patients undergoing surgical resection. The entire study was approved under a protocol approved by the University of Texas Southwestern Medical Center and Partners Institutional Review Board (IRB) (STU102010-051). All animal experiments were conducted under the Institutional Animal Care and Use Committee of the University of Texas Southwestern Medical Center approved guidelines for animal welfare (APN 2016-101380).

### 2.16. Data analysis

The statistical results were obtained from at least two independent biological replicates. Detailed n values for each panel in the figures are stated in the corresponding legends. A 2-tailed unpaired Student’s t-test was used to analyze data from experiments containing two groups. In comparison, one-way ANOVA was used to analyze data from experiments containing more than two groups. An appropriate post hoc test was used to analyze data that demonstrated significance in ANOVA. Kaplan-Meier (K-M) curves and log-rank tests were used for survival analysis. P☐<☐0.05 was considered to be statistically significant. Statistical analysis was performed using GraphPad Prism 9.4.1 software.

## 3. Results

### 3.1. CDK4/6i synergizes with PARPi in RB1-proficient human NSCLC cell lines

We first sought to establish the phenotypic implications of the combination of CDK4/6i (palbociclib) and PARPi (olaparib or rucaparib) through the use of clonogenic assays across a panel of NSCLC cell lines. Compared to monotherapy, combined use of palbociclib and olaparib (or palbociclib and rucaparib) induced a significantly stronger inhibitory effect on the clonogenic growth of RB-proficient NSCLC A549, H460, and H1437 cell lines (Fig. 1A and Supplementary Fig. 1A). To evaluate the nature of the combination effect between CDK4/6i and PARPi, we performed isobologram analysis using the CompuSyn software (19, 20). We calculated the combination indices (CIs) of various combinations of CDK4/6i and PARPi on A549, H460, and H1437 cells. A CI between 0 and 1 indicates a synergistic effect between two drugs, and the closer the index is to 0, the stronger the synergism (20). Combination indices ranged from 0.1 to 0.9 for all combinations. Notably, palbociclib showed strong synergism (CI < 0.9) with olaparib or rucaparib across a range of doses, indicating strong synergy and demonstrating that synergy was not limited to a specific dose combination. (Fig. 1B and Supplementary Fig. 1B).

**Figure 1.**
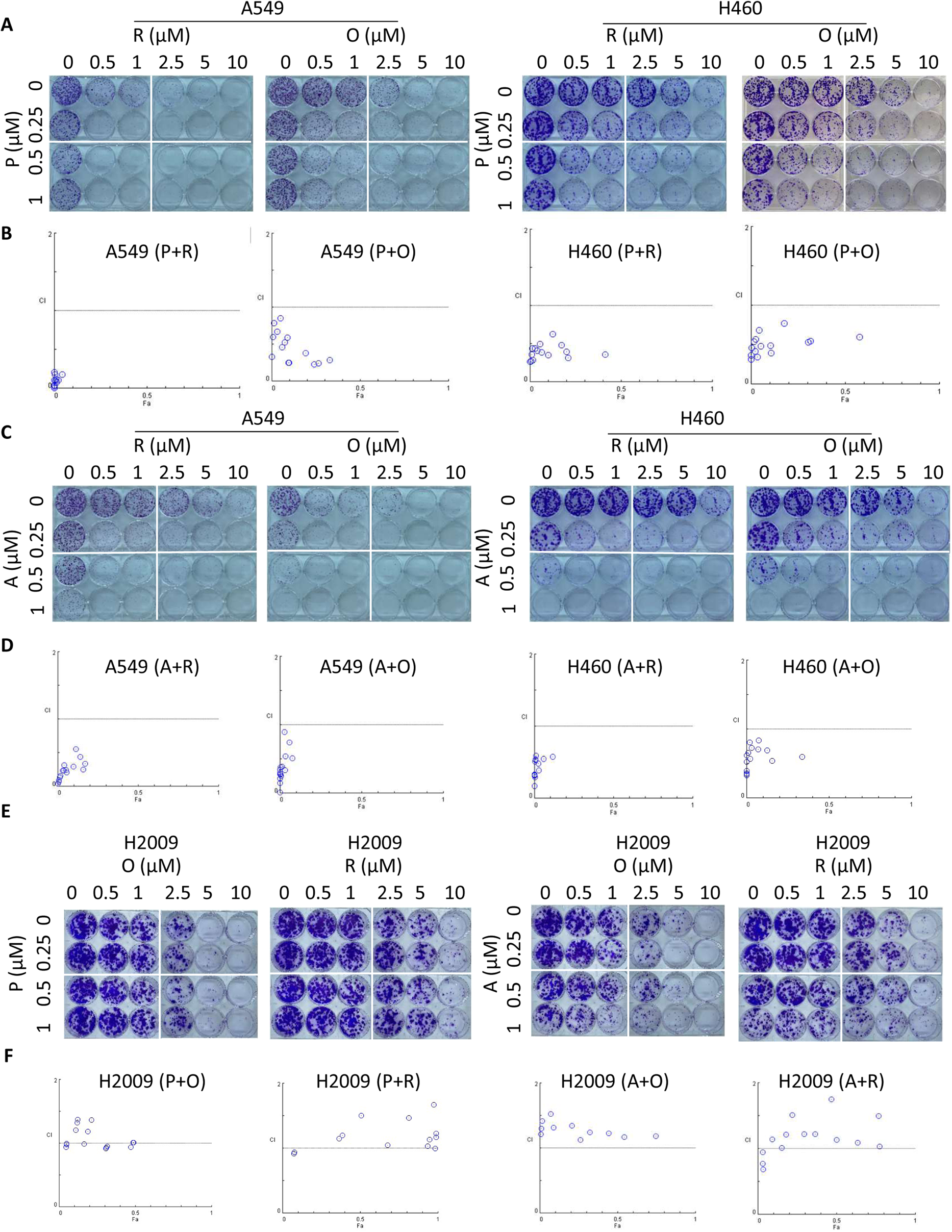
Combination treatment with CDK4/6i and PARPi produces synergistic inhibition of clonogenic survival in RB-proficient human NSCLC cell line models. (A) Colony formation assays in RB-proficient A549 and H460 NSCLC cell lines in the presence of vehicle (V, DMSO), or different concentrations of palbociclib (P), olaparib (O)/rucaparib (R), or combination (P+O/P+R) for 10-14 days. (B) Combination index (CI, Y-axis) values were calculated with CompuSyn software of P+O or P+R from the colony formation assay shown in A. The X-axis indicates the fraction affected (f a). All the CI values were less than 1, indicating synergy. (C) Colony formation assays in RB-proficient A549 and H460 NSCLC cell lines in the presence of vehicle (V, DMSO), or different concentrations of abemaciclib (A), olaparib (O)/rucaparib (R), or combination (A+O/A+R) for 10-15 days. (D) CI values were calculated with CompuSyn software of A+O/A+R from the colony formation assay shown in C. All the CI values were less than 1, indicating synergy. (E) Colony formation assays in RB-deficient H2009 NSCLC cell line in the presence of vehicle (V, DMSO), or different concentrations of palbociclib (P)/abemaciclib (A), olaparib (O)/rucaparib (R), or combination (A+O/A+R/P+O/P+R) for 10-14 days. (F) CI values calculated with CompuSyn software of P+O/P+R/A+O/A+R from colony formation assay shown in E. Most values were greater than 1, indicating no synergy in RB-deficient H2009 NSCLC cells.

We next performed clonogenic assays to examine whether PARPi (olaparib or rucaparib) could act synergistically with the structurally distinct CDK4/6i, abemaciclib, to inhibit clonogenic growth. Similarly, compared to single agents, we observed a superior inhibitory effect on clonogenic growth of RB-proficient NSCLC A549 and H460 cell lines when abemaciclib was used in combination with PARPi (olaparib or rucaparib) (Fig. 1C). Notably, combination indices ranged from 0.1 to 0.9, suggesting that the combination effect was synergistic (Fig. 1D). In contrast, no synergistic combination activity (CI>1) was observed in RB-deficient NSCLC H2009 cell line (Fig. 1E and F) and RB-deficient prostate cancer DU145 cell line (Supplementary Fig. 1C and D), suggesting that the observed synergy was dependent on the expression of RB protein in tumor cells.

We analyzed the growth-inhibitory effects of olaparib and palbociclib combination treatment using proliferation assays in A549 cells to further validate our findings. Two combination designs were used-constant drug ratio (1:1) and non-constant drug ratios. Compared with single-agent treatment, combining olaparib and palbociclib induced a significantly greater inhibitory effect on cell proliferation in both settings (Supplementary Fig. 1E and F). Combination indices ranged from 0.1 to 0.9 for all combinations, suggesting synergy in these two combination settings. Together, these data indicate that CDK4/6i synergized with PARPi in multiple RB-proficient NSCLC cell line models in different assay types and various combinations of these classes of agents.

### 3.2. CDK4/6i enhances PARPi-induced DNA damage in RB-proficient human NSCLC cells

We next assessed the molecular consequences of the synergistic interaction between CDK4/6i and PARPi. First, we sought to examine the effect of CDK4/6i and PARPi alone and in combination on cellular DNA damage. Immunofluorescence staining analysis revealed that, when compared with single-agent or vehicle treatment, the combination of palbociclib and olaparib induced significantly more phosphorylated histone H2AX (γH2AX) nuclear foci (a surrogate marker for DSBs) after 72 h treatment. In contrast to either agent alone, the combination of palbociclib and olaparib resulted in a marked increase in the number of cancer cells with high-intensity, pan-nuclear γH2AX staining as well as the total number of γH2AX positive cancer cells (Fig. 2A). These findings were validated by immunoblot analysis showing that the protein level of γH2AX was also significantly increased by the combination treatment of palbociclib and olaparib (Fig. 2B). We next examined whether CDK4/6i would increase PARPi-induced DNA damage by using alkaline comet assay, which detects and measures both DNA SSBs and DSBs within cells. Again, the combination treatment of palbociclib and olaparib induced a significant accumulation of damaged DNA in A549 (Fig. 2C) and H460 cells (Supplementary Fig. 2A), in contrast to either agent alone. Since palbociclib and olaparib can arrest cells in different cell cycle phases, we next analyzed their effect on cell-cycle progression. As expected, flow cytometry analysis revealed that single-agent treatment by palbociclib arrested NSCLC cells in the G1 phase. In contrast, treatment with olaparib arrested cells in the G2-M phase (Fig. 2D). Interestingly, the combined use of olaparib and palbociclib increased the percentage of cells arrested in the G1 phase and decreased the percentage of cells arrested in the S phase to a greater extent than either single agent alone in A549 (Fig. 2D) and H460 cells (Supplementary Fig. 2B).

**Figure 2.**
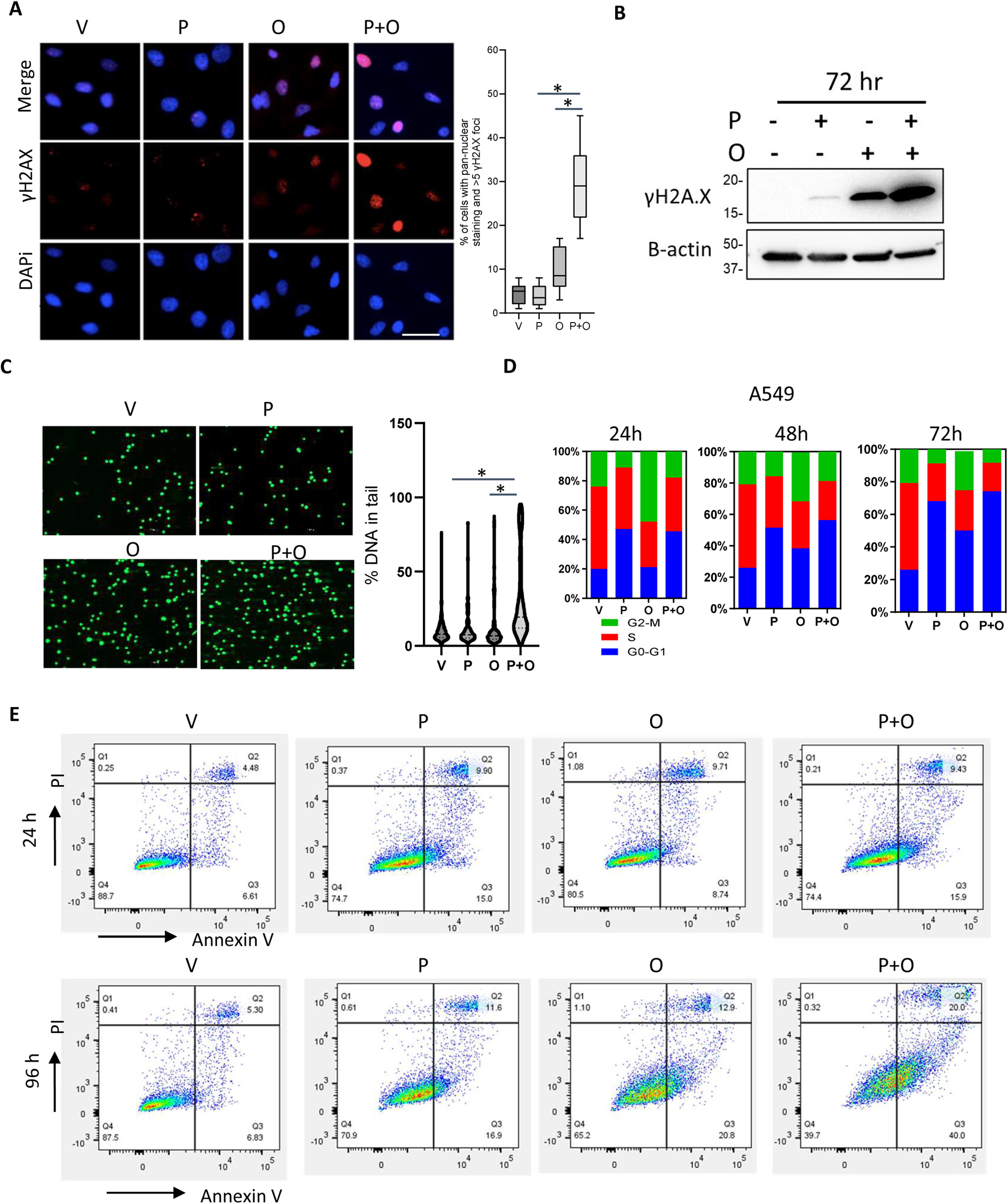
CDK4/6i increases PARPi-induced DNA damage in RB-proficient human NSCLC cells. A549 cells were treated with vehicle (V, DMSO), palbociclib (P, 1μM), and olaparib (O, 2 μM) alone or in combination (P+O) for 72 h. (A) DNA damage was measured by immunofluorescence microscopy with anti-γH2AX antibody after 72 h treatments with V, P, O, or P+O. Left, representative images of each treatment and right plot, quantitation of the percentage of cells with either pan-nuclear staining or > 5 foci. Scale bar, 40 μm. (B) Western blot detection of γH2AX in A549 cells after 72 h treatments with V, P, O, or P+O as above. (C) Alkaline comet assay of A549 cells treated with V, P, O, and P+O for 72 h. Representative images of the quantification of the alkaline comet assay (left) and the tail moment (right) were shown. (D) Flow cytometry analysis of cell cycle distribution in A549 cells treated with V, P, O, or P+O for the indicated time. (E) Annexin V/PI double staining apoptosis analysis by flow cytometry in A549 cells treated with V, P, O, or P+O for the indicated time. Box and whisker plots represent values within the interquartile range (boxes) and the minimum to maximum (whiskers). The line within the box shows the median. *P☐<☐0.01 *vs.* single agent treatment, by 1-way ANOVA with Tukey’s multiple comparisons test.

Further, we utilized flow cytometry to measure the induction of apoptosis. Consistent with drug-induced clonogenic growth inhibition effect, combined use of olaparib and palbociclib but not either agent alone induced substantial apoptotic cell death as determined by percent positivity of annexin V/PI staining (Fig. 2E and Supplementary Fig. 2C). Taken together, these results indicate that, compared with vehicle or single-agent treatment, olaparib and palbociclib combination treatment increase DNA damage, interrupt cell-cycle checkpoints, and enhance apoptotic cell death in RB-proficient NSCLC cells.

### 3.3. CDK4/6i promotes PARP1 protein degradation in RB-proficient NSCLC cells

Given that PARP1 is a critical DNA damage sensor, it recruits downstream proteins by poly ADP-ribosylation (PARylation) upon activation (23). We analyzed the protein levels of PARP1 after CDK4/6i (palbociclib or abemaciclib) treatment in RB-proficient NSCLC cell lines (A549 and H1437) by immunoblot analysis. Interestingly, the protein levels of PARP1 drastically decreased upon palbociclib (or abemaciclib) treatment in a time-and dose-dependent manner in RB-proficient NSCLC cell lines (Fig. 3A, 3B, and Supplementary Fig. 3A), whereas similar observations were not seen in the RB-deficient H2009 cells (Supplementary Fig. 3A). Our data showed that abemaciclib effectively reduced PARP1 protein level within 2-8 h and achieved near-complete PARP1 depletion within 24 to 48 h (Supplementary Fig. 3C). Additionally, the decrease of PARP1 protein level by CDK4/6i was associated with reduction of PARP1 catalytic activity (inhibition of PARylation) (Fig. 3C). Interestingly, CDK4/6i did not significantly decrease PARP1 mRNA levels within 24 h treatment (Supplementary Fig. 3B) in A549 and H1437 cells, suggesting that CDK4/6i regulates PARP1 mainly at the protein level rather than at the gene expression level. Consistent with this idea, we observed that CDK4/6i-induced PARP1 degradation was inhibited by co-treatment with the proteasome inhibitor MG132 (Fig. 3C), suggesting that CDK4/6i induced PARP1 degradation in a proteasome-dependent manner. We also observed that CDK4/6i (palbociclib or abemaciclib) significantly decreased the half-life of endogenous PARP1 protein in the presence of protein synthesis inhibitor, cycloheximide (CHX) (Fig. 3D). To determine if PARP1 downregulation by CDK4/6i is a general feature of cells arrested in G1 phase of the cell cycle, we assessed the PARP1 protein levels in A549 cells in the presence of dexamethasone, which induces cell cycle arrested in G0/G1 (24). Strikingly, PARP1 protein degradation was not observed in cells arrested in the G1 phase by dexamethasone (Supplementary Fig. 3D), suggesting that PARP1 downregulation is not a general feature of cells arrested in the G1 phase. Since most DNA damage-induced PARylation is primarily mediated by PARP1 (25), we examined the effect of CDK4/6i on DNA damage-induced PARylation. We treated A549 cells with CDK4/6i followed by exposure to methyl methanesulfonate (MMS), an alkylating agent that induces DNA damage. PAR levels were reduced by palbociclib and abemaciclib treatment even in the presence of 0.01% MMS (Fig. 3E), suggesting that CDK4/6i blocked DNA damage-induced PARylation. These results reveal that CDK4/6i negatively regulates PARP1 protein stability, substantially impacting the PARylation function.

**Figure 3.**
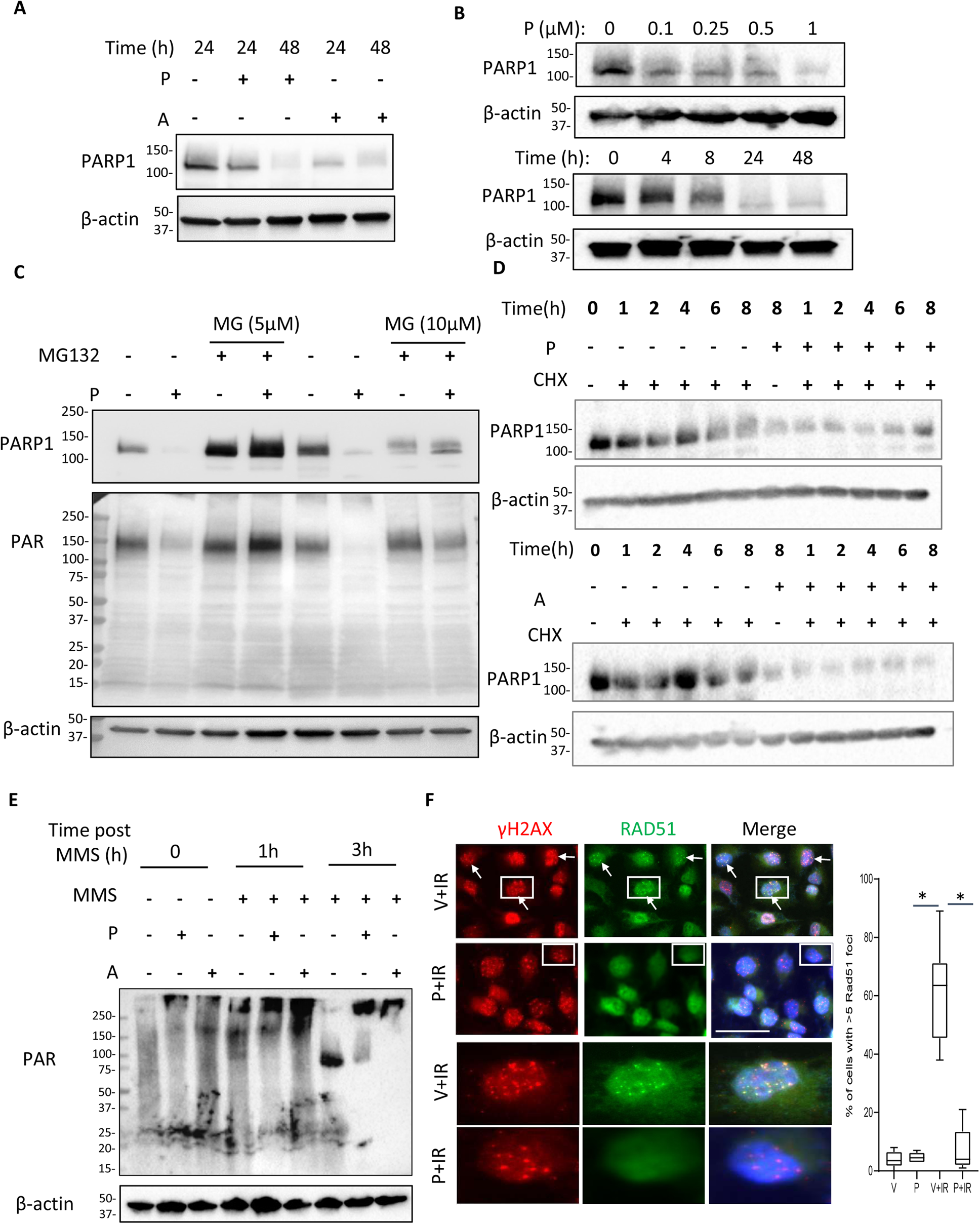
CDK4/6is promote PARP1 protein degradation and regulate DNA repair factor availability and DNA repair competency in RB-proficient NSCLC cells. (A) Western blot detection of PARP1 levels in A549 cells treated with palbociclib (P, 1 µM) or abemaciclib (A, 1 µM) for the indicated times. (B) Western blot detection of PARP1 levels in A549 cells treated with increasing concentrations of palbociclib (P) for 24 h (top) and with P (1 µM) for the indicated times (bottom). (C) Co-treatment with the proteasome inhibitor MG132 inhibits CDK4/6i-induced PARP1 degradation. A549 cells were treated with palbociclib (1 µM) for 16 h and MG132 for an additional 6 h. PARP1 and PAR in total cell lysates were analyzed by Western blot. (D) Western blot detection of PARP1 levels in A549 cells treated with control or CDK4/6i (palbociclib or abemaciclib, 1☐μM) in the presence of cycloheximide (CHX; 10☐μg☐/ml) for the indicated times. (E) Western blot detection of PARylation levels in A549 cells treated with 1 μM palbociclib or abemaciclib for 2 h, followed by MMS (0.01%) for an additional 1 or 3 h. (F) Representative images (left) of γH2AX (red) and RAD51 (green) staining in A549 cells treated with vehicle (V, DMSO), palbociclib (P, 1 μM) for 16 h and with/without IR (2Gy) for additional 4 h. Cells with white squares are enlarged below for a better visualization. The arrows indicate the position of foci. The box and whisker plot (right) quantifies cells with more than 5 foci. *P☐<☐0.01 V+IR *vs*. P+IR, by 1-way ANOVA with Tukey’s multiple comparisons test. Scale bar, 40 μm.

### 3.4. CDK4/6i regulates DNA repair factors availability and confers sensitivity to radiation and bleomycin

It is well established that PARylation occurs at or near the sites of DNA damage and promotes the recruitment of DNA repair factors via their PAR binding domains (23). Given the robust effect of CDK4/6i on PARP1 protein stability and PARylation, we assessed the impact of CDK4/6i treatment on IR-induced recruitment of RAD51 nuclear foci as a surrogate HR functionality marker and phosphorylated DNA-dependent protein kinase catalytic subunit (pDNA PKcs) and 53BP1 nuclear foci, as surrogate NHEJ functionality markers. IR (2Gy) treatment increased RAD51 foci in∼70 % of cells (P < 0.01) compared to no IR control (∼3 %), indicative of an accumulation of DSBs requiring HR-mediated resolution (Fig. 3F). In contrast, pretreatment with CDK4/6i (palbociclib, 1µM) did not produce an increase in IR-induced RAD51 foci, despite the presence of DNA damage as indicated by γH2AX foci. CDK4/6i had no significant effect on IR-induced NHEJ markers (pDNA PKcs and 53BP1) (Supplementary Fig. 3E and F). These data suggest the absence of a robust HR response in the presence of CDK4/6i. Given the observed effect of CDK4/6i on the recruitment of DNA repair factors, we tested the impact of CDK4/6i on sensitivity to DNA-damaging agents such as IR and bleomycin, commonly used in clinics to treat NSCLC patients. Two NSCLC cell lines were treated with palbociclib (0.5 and 1 µmol/L) for 2 h and then exposed to increasing radiation dosage intervals (0, 1, 2, and 3 Gy), and the clonogenic growing effects were analyzed. Treatment with palbociclib led to a significant radiosensitization effect in the A549 and H460 cell lines (Supplementary Fig. 3G). In addition, concomitant therapy with palbociclib and bleomycin also resulted in a synergistic inhibitory effect on the clonogenic growth of A549 cells (Supplementary Fig. 3H). These results reveal that CDK4/6i decreases the availability of DNA repair factors involved in HR and confers sensitivity to DNA-damaging agents, such as radiation and bleomycin.

### 3.5. PARP trapping is essential for the synergy between CDK4/6i and PARPi

To further understand the molecular effects elicited by combined PARP and CDK4/6 inhibition, we first looked at the PARP1 protein levels and the PARylation status under the drug combination. Similar to CDK4/6i alone, PARP1 protein levels were reduced by the combination of CDK4/6i and PARPi. Interestingly, the reduction in PARylation was more significant upon palbociclib and olaparib combination treatment than vehicle or single-agent treatment. Consistent with previous results, we observed these reductions in PARP1 and PARylation in RB-proficient A549 cells but not in RB-deficient H2009 cells (Fig. 4A).

**Figure 4.**
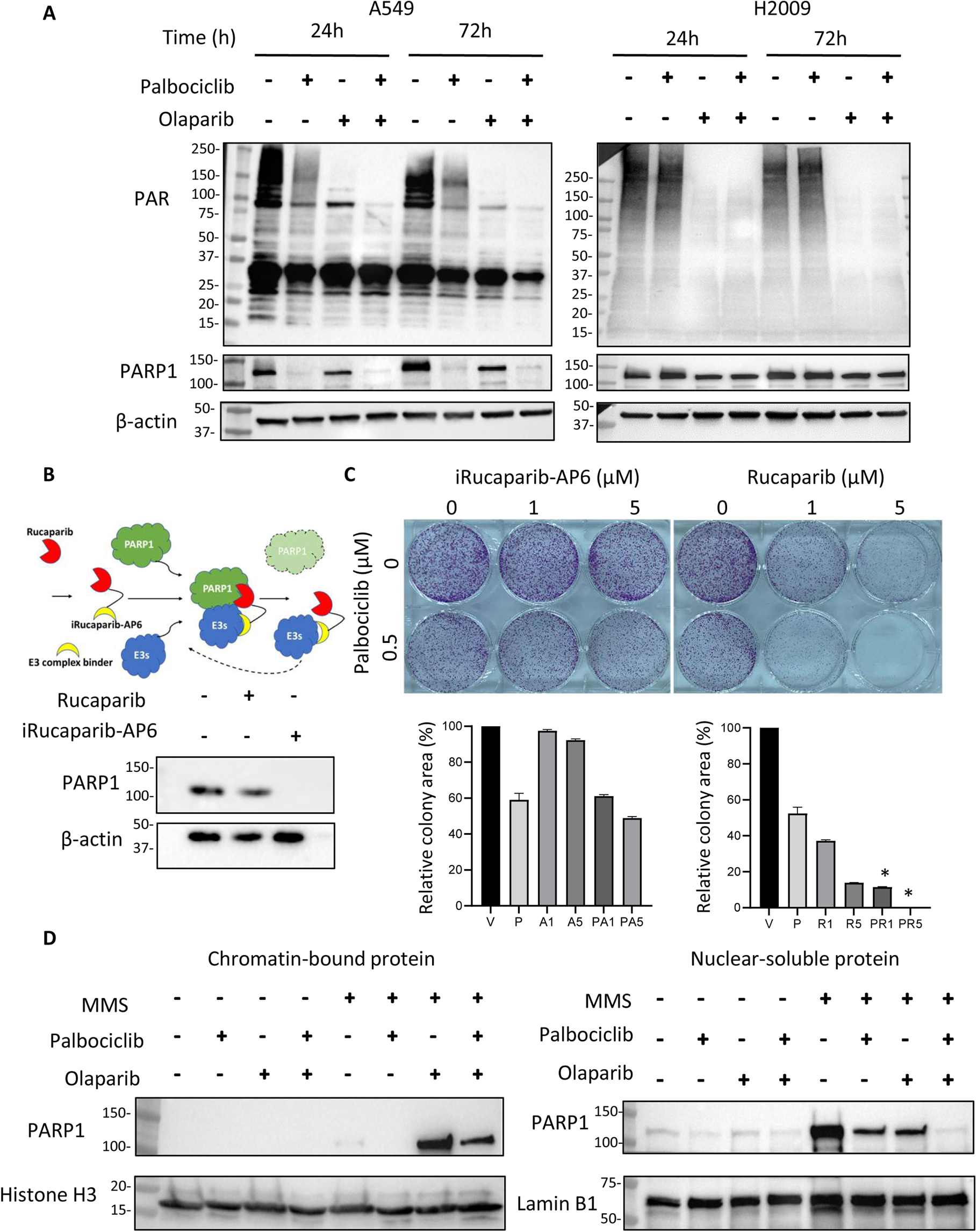
PARP trapping is essential for the synergy between CDK4/6i and PARPi. (A) Western blot analysis of whole cell lysates obtained from the A549 and H2009 cells after 24 and 72 h of treatment with vehicle (V, DMSO), palbociclib (P, 1 μM), olaparib (O, 2 μM) or combination (P+O). Blots were probed with the indicated antibodies, and β-actin was used as a loading control. (B) Schematic of the iRucaprib-AP6 mechanism (top) and western blot showing the effect of iRucaprib-AP6 on PARP1 degradation (bottom). A549 cells were treated with rucaparib (1☐μM) or iRucaparib-AP6 (1☐μM) for 48☐h. Total proteins were extracted and analyzed using the indicated antibodies. (C) Representative images of colony forming assays (top) and quantification (bottom) in A549 cells in the presence of vehicle (V, DMSO), palbociclib (P, 0.5 μM), rucaparib (R, 1 and 5 μM)/iRucaparib-AP6 (A, 1 and 5 μM), or combination (PA1, PA5, PR1, PR5). *P☐<☐0.001 *vs.* single agent treatment, by 1-way ANOVA with Tukey’s multiple comparisons test. (D) Western blot analysis of chromatin-bound and nuclear-soluble fractions prepared from A549 cells. Cells were pretreated with vehicle (DMSO), palbociclib (1☐μM), olaparib (2☐μM) or combo for 16 h followed by a 1 h treatment with MMS (0.01%). Chromatin-bound and nuclear-soluble proteins were extracted and analyzed using the indicated antibodies. Histone H3 and Lamin B1 were used as chromatin-bound and nuclear-soluble fractions markers, respectively.

Given the critical role of PARP1 trapping in the PARPi-induced cytotoxicity (26), we tested the relative contributions of PARP1 trapping and PARP catalytic inhibition in the synergy between CDK4/6i and PARPi. iRucaparib-AP6 is a derivative of rucaparib with a highly efficient and specific PARP1 degradation activity based on the Proteolysis Targeting Chimera (PROTAC) approach (17). iRucaparib-AP6 recruits the E3 complex to PARP1 to induce its ubiquitination and subsequent proteasomal degradation (Fig.4B, top) (17). Both iRucaparib-AP6 and rucaparib have similar PARP catalytic inhibition capacities (17). Contrarily to the parent rucaparib, iRucaparib-AP6 led to robust downregulation of PARP1 in A549 cells (Fig. 4B, bottom). We found that treatment of A549 cells with palbociclib in combination with rucaparib, but not with iRucaparib-AP6, significantly increased the clonogenic growth inhibition effect (Fig. 4C), suggesting that PARP trapping is essential for the synergy between PARPi and CDK4/6i. To further distinguish the effects of PARP catalytic inhibition from PARP1 trapping on the synergy, we used veliparib, another PARP inhibitor with less potent PARP1 trapping capacity than olaparib and rucaparib (26). Treatment with veliparib alone produced a limited clonogenic growth inhibition effect and did not significantly potentiate palbociclib-mediated inhibition in A549 (Supplementary Fig. 4 A and B). The finding that neither iRucaparib-AP6 nor veliparib synergized when given in combination with CDK4/6i suggests that inhibition of PARP1 catalytic activity is insufficient to produce synergy when combined with CDK4/6 inhibition. These findings contrast with the significant increase in clonogenic growth inhibition caused by olaparib or rucaparib when combined with palbociclib or abemaciclib. These data support the importance of PARP1 trapping as a mechanism of synergy by combining CDK4/6i and PARPi. To determine whether CDK4/6i could enhance PARP1 trapping, we used a sub-cellular fractionation assay to assess nuclear and chromatin-bound levels of PARP1. Consistent with previous reports in the literature, we could not detect an increase in PARP1-DNA complexes with PARPi (olaparib) alone, likely due to the lack of sensitivity of the trapping assay (26). In the presence of MMS, olaparib, but not palbociclib, greatly induced PARP1 accumulation in the chromatin-bound fraction (Fig. 4D), and the combination of both compounds did not increase this accumulation above that induced by the PARPi alone. Since CDK4/6i did not further enhance the chromatin-bound fraction of PARP1, we conclude that the synergistic action between CDK4/6i and PARPi cannot be explained by elevated PARP1 trapping-induced cytotoxicity alone. Since CDK4/6i promotes the degradation of PARP1 protein, we hypothesized that CDK4/6i could also impede the DNA damage response *via* a decreased basal level of PARP1. To test this hypothesis, we assayed the impact of the combination treatment of palbociclib and olaparib on IR-induced BRCA1 and 53BP1 foci formation in NSCLC A549 cells (Supplementary Fig. 4 C). BRCA1 and 53BP1 are determinants of the choice between HR and NHEJ for DNA damage repair (27, 28);. At the same time, BRCA1 promotes HR, and 53BP1 stimulates NHEJ. In response to DNA damage, A549 cells displayed well-characterized BRCA1 foci formation, even in the presence of CDK4/6i (palbociclib) or PARPi (olaparib) alone. Still, this phenotype was not observed in the combination of CDk4/6i and PARPi (Supplementary Fig. 4 C). On the other hand, cells treated with CDK4/6i and PARPi combination exhibited a similar 53BP1 foci staining compared to control or single agents treated cells (Supplementary Fig. 4 C). These data suggest the absence of a robust HR response in the presence of CDK4/6i and PARPi. We further examined the potentiating capability of the combination of palbociclib and olaparib on DNA damage repair by combining low-dose of palbociclib (100 and 200 nM) and Olaparib (100 and 200 nM) with IR. Compared with single-agent treatment, we observed a significant increase in sensitivity to IR in A549 cells treated with low dose palbociclib and olaparib combination (Supplementary Fig. 4 D). Taken together, these data show that the combination of PARPi and CDK4/6i inhibits DDR and PARP trapping is essential for the synergistic combination.

### 3.6. The combination of CDK4/6i (Palbociclib) and PARPi (olaparib) is well tolerated and has superior survival benefits in mice

To assess the therapeutic effect of the CDK4/6i and PARPi combination *in vivo*, mice bearing A549 tumor xenografts s.c. were treated with either palbociclib (dosed once daily by oral gavage, 100☐mg/kg), olaparib (dosed once daily by intraperitoneal injection, 50☐mg/kg), or the combination of palbociclib and olaparib for 21 days. The tumors in the vehicle group proliferated to the maximum size threshold within 21 days, recapitulating the aggressive nature of NSCLC in human patients. While a single treatment of olaparib or palbociclib only slightly slowed tumor growth, the combination treatment of olaparib and palbociclib led to stable disease in most tumors, and tumor size remained stable throughout treatment. During the treatment period, all animals treated with vehicle or single agents had to be euthanized except two in the palbociclib-alone group; however, no animals died in the presence of the combination, suggesting the combination of palbociclib and olaparib significantly slowed down tumor growth and led to substantial survival benefit (Fig. 5A and B). Notably, the combination treatment of palbociclib and olaparib was well tolerated as no significant change in body weight was observed during the study (Fig. 5C). In addition, as seen *in vitro*, palbociclib and olaparib combination treatment profoundly reduced cell proliferation *in vivo*, as measured by immunohistochemical staining for Ki67in the A549 xenografts (Fig. 5D), suggesting that these agents in combination induce a complete cell cycle arrest compared with each agent alone. Finally, microscopic examination of hematoxylin-eosin-stained tissue sections of vital organs (liver, heart, and kidney) did not show any significant histological change that would indicate toxic effects of palbociclib and olaparib, suggesting the combination of palbociclib and olaparib was well tolerated in mice.

**Figure 5.**
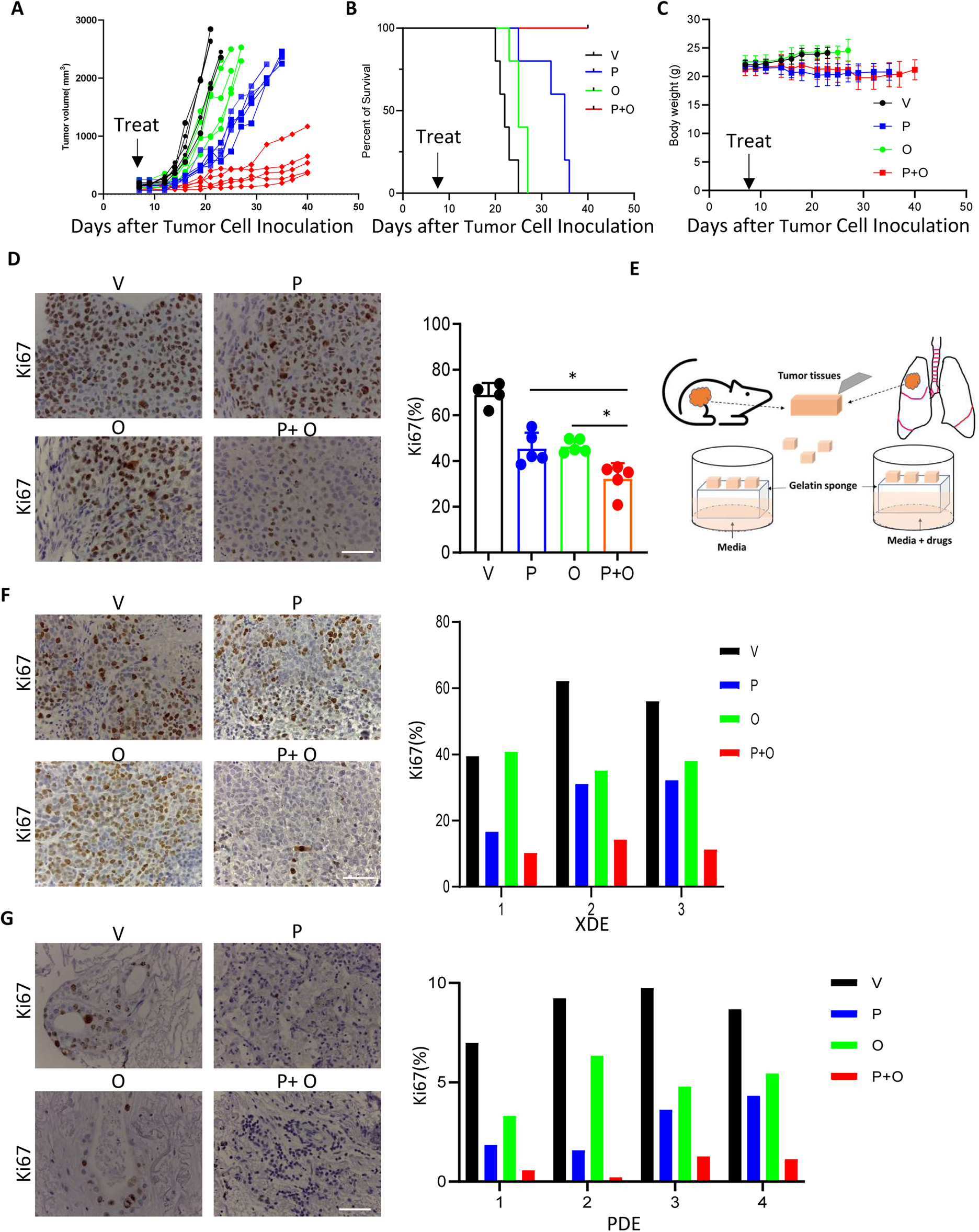
Combined PARP and CDK4/6 inhibition demonstrates superior antitumor efficacy against human NSCLC cell line xenografts, xenografts-derived explants (XDE), and patient-derived explants (PDE) models. (A) Tumor growth of A549 xenografts subjected to the indicated treatments: vehicle (V), palbociclib (P, 100 mg/kg daily), olaparib (O, 50 mg/kg daily), or the combination (P+O) for 21 days. n = 5 per group. (B) Cumulative survival of mice before tumor outgrowth to 2,500 mm^3^ vs. time. The Kaplan-Meier survival curve was plotted using GraphPad software. N=5 mice per group and the trend in survival is compared between respective vehicle and drug-treated groups (Log-rank test p<0.001). (C) Mice body weights were measured and are expressed as mean ± SD. (D) Representative images of immunodetection of the cell proliferation marker Ki67 in tumor sections from the A549 xenografts subjected to the indicated treatments for 14 days. The histogram shows the quantification of Ki67-positive cells. Data represent the mean ± SD, n = 4-5 per group. *P < 0.05, vs. single agent treatment, by 1-way ANOVA with Tukey’s multiple comparisons test. (E) Schematic illustration of tumor-derived explant models. Following surgical resection, xenografts or primary NSCLC tumors were cultured *ex vivo* with either vehicle or different treatments, as shown by the schematic illustration of the XDE and PDEs experiment. (F) Representative images of immunodetection of the cell proliferation marker Ki67 in tumor sections from xenografts-derived explants treated with V (DMSO), P (1☐μM), O (2☐μM) and P+O (1☐μM+2☐μM) for 72 h. The histogram shows the quantification of Ki67-positive cells in individual XDE. (G) Representative images of immunodetection of the cell proliferation marker Ki67 in tumor sections from PDEs treated with V (DMSO), P (1☐μM), O (2☐μM) and P+O (1☐μM+2☐μM) for 72 h. The histogram shows the quantification of Ki67-positive cells in individual PDE. Scale bars: 50 μm.

### 3.7. Combining CDK4/6i and PARPi decreases cell proliferation in xenografts-derived explants (XDE) and patient-derived explants (PDE)

We further validated the antitumor effect of combining CDK4/6i and PARPi on well-established tumor-derived explant models (Fig. 5E). The combination of palbociclib and olaparib, as determined by histological analysis, caused a substantial reduction in Ki67-positive A549 XDE cells (Fig. 5F). Thus, decreased proliferation may, at least in part, explain the superior treatment response to the combined PARPi-CDK4/6i regimen. To determine whether the combination of the CDK4/6i and PARPi can be applied in the clinic, we used NSCLC PDE models, which faithfully resemble the original characteristics of primary tumors, such as heterogeneity and histological structures. As determined by the histological analysis, the combination treatment caused substantially reduced Ki67 staining positive cells (Fig. 5G). Thus, combining CDK4/6i and PARPi is an effective therapy for NSCLC in XDE and PDE models. Together, these data open the possibility to the potential clinical utility of CDK4/6i and PARPi combination in treating NSCLC.

## 4. Discussion

In this preclinical study, we demonstrated that CDK4/6i (palbociclib and abemaciclib) synergizes with PARPi (olaparib and rucaparib) in the setting of RB-proficient human NSCLC models. We further showed that PARP1 trapping is required for the synergy between CDK4/6i and PARPi. Our data suggest that CDK4/6is promote PARP1 protein degradation, which impedes the ability to repair DNA damage. We further established that this repair deficiency could induce synthetic lethality when deployed in combination with PARPi with or without radiation and other DNA damage agents, such as bleomycin.

Most clinical trials that investigated CDK4/6i either as a single agent or in combination with different chemotherapy regimens in different treatment lines showed minimal activity, likely due to the early occurrence of acquired resistance and the lack of adequate biomarker selection. NSCLC is characterized by multiple mutations, including in the growth suppressor gene CDKN2A. Indeed, the loss of function of CDKN2A may account for up to 40% of lung adenocarcinomas (5). In addition, the clinical relevance of our findings stems from the fact that loss of pRB protein expression is only found in 15-30% of NSCLC patients (29, 30), suggesting that most patients with NSCLC could benefit from combining CDK4/6i and PARPi. CDK4/6i has previously been found to favor NHEJ while inhibiting HR, although when used alone, this shift did not alter the overall rate of DNA repair (31). Our data reveal, for the first time, that CDK4/6i may significantly impact DDR by regulating the protein stability of a critical component of DDR, PARP1. Moreover, the observed potentiating effect of low-dose olaparib and palbociclib combination on sensitivity to IR suggests that low-dose drug combinations might open an avenue for further clinical evaluation.

The mechanism for the CDK4/6i induced PARP1 degradation is an important open question that warrants further investigation because PARP1 is a multifunctional nuclear protein that participates in numerous critical cellular processes, including DNA damage, chromatin regulation, gene expression, ribosome biogenesis, and RNA biogenesis and metabolism (32, 33). CDK4/6i has previously been found to repress PARP1 transcription, and it is speculated to act through the induction of a repressive complex containing RB at the PARP1 promoter (34). In this sense, we also detected a moderate effect in PARP1 mRNA levels at 24 h. Still, we found that CDK4/6i has a powerful effect inducing PARP1 protein degradation with a high-speed kinetics that becomes evident at around 2 h after drug treatment. We also noted that the effect of CDK4/6i on PARP1 degradation is not merely a consequence of cell arrest in G1 but rather a specific effect of CDK4/6i. PARP1 trapping is a crucial driver of PARPi-induced bone marrow toxicity, which may be necessary for combination regimens, including DNA-damaging chemotherapy that also causes bone marrow toxicity (35). Our data reveal that combining CDK4/6i with PARPi reduces the overall level of PARP1, thus the trapping potency of PARPi, which could lead to less toxicity. Of note, recent preclinical data indicate that CDK4/6i protects against radiation-induced intestinal injury in mice by inducing protective DDR selectively in normal tissues with WT p53, but not in cancers with deregulated p53 (36)Future studies are necessary to evaluate the effect of the combination CDK4/6i and PARPi in non-transformed cells and identify possible side effects associated with it, with or without radiotherapy or other DNA damage agents.

Given that these therapeutic modalities are already available for clinical use, our approach of combining PARPi with CDK4/6i represents an accessible treatment strategy for NSCLC patients. Our findings could open new research horizons, as therapeutic applications of PARPi will not be limited to HR-deficient populations. Importantly, we could identify a biomarker-set to guide patient selection for this combination strategy. Because CDK4/6i rely on the phosphorylation and inactivation of RB and loss of RB is considered as a marker of resistance to CDK4/6 inhibition (37), patients could be screened for their RB status, followed by selection of CDK4/6i to induce PARP1 protein degradation in RB-proficient tumors. This could significantly lead the current clinical practice for treating NSCLC towards a more precision medicine-based approach and offer NSCLC patients the best possible outcomes. Overall, our data expand our understanding of the DDR consequences of CDK4/6 inhibition in NSCLC and support the feasibility of combining CDK4/6i and PARPi in NSCLC patients with RB-proficient tumors.

## Acknowledgments

This work was supported by Congressionally Directed Medical Research Programs LCRP under award number W81XWH-19-1-0645 and the National Cancer Institute of the National Institutes of Health under award number 5 P30 CA142543 09. We thank Dr. John D. Minna (The University of Texas Southwestern Medical Center, Dallas, TX) for sharing human NSCLC cell lines and Dr. Yonghao Yu (The University of Texas Southwestern Medical Center, Dallas, TX) for sharing iRucaparib-AP6.

## CRediT authorship contribution statement

**Carlos M Roggero**: Writing – review & editing, Writing – original draft, Validation, Investigation, Formal analysis, Data curation, Conceptualization.

**Anwesha B Ghosh:** Writing – review & editing, Writing – original draft, Validation, Investigation, Formal analysis, Data curation, Conceptualization.

**Anvita Devineni:** Writing – review & editing, Data curation.

**Shihong Ma:** Writing – review & editing, Data curation, Validation, Methodology, Investigation,

**Eliot Blatt:** Writing – review & editing, Data curation, Methodology, Investigation,

**Ganesh V. Raj:** Writing – review & editing, Resources, Supervision, Funding acquisition.

**Yi Yin:** Writing – review & editing, Writing – original draft, Investigation, Formal analysis, Data curation, Validation, Supervision, Resources, Investigation, Conceptualization, Funding acquisition.

## Declaration of competing interest

G.V.R. holds issued and pending patents, which have been licensed to EtiraRx. G.V.R. serves or has served in an advisory role to Bayer, Johnson and Johnson, Myovant, EtiraRx, Amgen, Pfizer, and Astellas. He has or has had grant support from Bayer, EtiraRx, and Johnson and Johnson. All other authors declare that they have no competing interests.

## References

1. Esposito V, Baldi A, Tonini G, Vincenzi B, Santini M, Ambrogi V, et al. Analysis of cell cycle regulator proteins in non-small cell lung cancer. J Clin Pathol. 2004;57(1):58–63.

2. Eymin B, and Gazzeri S. Role of cell cycle regulators in lung carcinogenesis. Cell Adh Migr. 2010;4(1):114–23.

3. Blain SW. Switching cyclin D-Cdk4 kinase activity on and off. Cell Cycle. 2008;7(7):892–8.

4. Cancer Genome Atlas Research N. Comprehensive genomic characterization of squamous cell lung cancers. Nature. 2012;489(7417):519-25.

5. Cancer Genome Atlas Research N. Comprehensive molecular profiling of lung adenocarcinoma. Nature. 2014;511(7511):543-50.

6. Sherr CJ, Beach D, and Shapiro GI. Targeting CDK4 and CDK6: From Discovery to Therapy. Cancer Discov. 2016;6(4):353–67.

7. Edelman MJ, Redman MW, Albain KS, McGary EC, Rafique NM, Petro D, et al. SWOG S1400C (NCT02154490)-A Phase II Study of Palbociclib for Previously Treated Cell Cycle Gene Alteration-Positive Patients with Stage IV Squamous Cell Lung Cancer (Lung-MAP Substudy). J Thorac Oncol. 2019;14(10):1853-9.

8. Zhang J, Xu D, Zhou Y, Zhu Z, and Yang X. Mechanisms and Implications of CDK4/6 Inhibitors for the Treatment of NSCLC. Front Oncol. 2021;11:676041.

9. Solomon B, Callejo A, Bar J, Berchem G, Bazhenova L, Saintigny P, et al. A WIN Consortium phase I study exploring avelumab, palbociclib, and axitinib in advanced non-small cell lung cancer. Cancer Med. 2022;11(14):2790–800.

10. Khanna KK, and Jackson SP. DNA double-strand breaks: signaling, repair and the cancer connection. Nat Genet. 2001;27(3):247–54.

11. Juncheng P, Joseph A, Lafarge A, Martins I, Obrist F, Pol J, et al. Cancer cell-autonomous overactivation of PARP1 compromises immunosurveillance in non-small cell lung cancer. J Immunother Cancer. 2022;10(6).

12. Michels J, Adam J, Goubar A, Obrist F, Damotte D, Robin A, et al. Negative prognostic value of high levels of intracellular poly(ADP-ribose) in non-small cell lung cancer. Ann Oncol. 2015;26(12):2470–7.

13. Remon J, Besse B, Leary A, Bieche I, Job B, Lacroix L, et al. Somatic and Germline BRCA 1 and 2 Mutations in Advanced NSCLC From the SAFIR02-Lung Trial. JTO Clin Res Rep. 2020;1(3):100068.

14. Passiglia F, Reale ML, Cetoretta V, Parlagreco E, Jacobs F, Listi A, et al. Repositioning PARP inhibitors in the treatment of thoracic malignancies. Cancer Treat Rev. 2021;99:102256.

15. Yang Y, Luo J, Chen X, Yang Z, Mei X, Ma J, et al. CDK4/6 inhibitors: a novel strategy for tumor radiosensitization. J Exp Clin Cancer Res. 2020;39(1):188.

16. Naz S, Sowers A, Choudhuri R, Wissler M, Gamson J, Mathias A, et al. Abemaciclib, a Selective CDK4/6 Inhibitor, Enhances the Radiosensitivity of Non-Small Cell Lung Cancer In Vitro and In Vivo. Clin Cancer Res. 2018;24(16):3994–4005.

17. Wang S, Han L, Han J, Li P, Ding Q, Zhang QJ, et al. Uncoupling of PARP1 trapping and inhibition using selective PARP1 degradation. Nat Chem Biol. 2019;15(12):1223–31.

18. Gilbreath C, Ma S, Yu L, Sonavane R, Roggero CM, Devineni A, et al. Dynamic differences between DNA damage repair responses in primary tumors and cell lines. Transl Oncol. 2021;14(1):100898.

19. Chou TC. Drug combination studies and their synergy quantification using the Chou-Talalay method. Cancer Res. 2010;70(2):440–6.

20. Chou TC, and Talalay P. Quantitative analysis of dose-effect relationships: the combined effects of multiple drugs or enzyme inhibitors. Adv Enzyme Regul. 1984;22:27–55.

21. Yin Y, Li R, Xu K, Ding S, Li J, Baek G, et al. Androgen Receptor Variants Mediate DNA Repair after Prostate Cancer Irradiation. Cancer Res. 2017;77(18):4745–54.

22. Gyori BM, Venkatachalam G, Thiagarajan PS, Hsu D, and Clement MV. OpenComet: an automated tool for comet assay image analysis. Redox Biol. 2014;2:457–65.

23. Gupte R, Liu Z, and Kraus WL. PARPs and ADP-ribosylation: recent advances linking molecular functions to biological outcomes. Genes Dev. 2017;31(2):101–26.

24. Greenberg AK, Hu J, Basu S, Hay J, Reibman J, Yie TA, et al. Glucocorticoids inhibit lung cancer cell growth through both the extracellular signal-related kinase pathway and cell cycle regulators. Am J Respir Cell Mol Biol. 2002;27(3):320–8.

25. Beck C, Robert I, Reina-San-Martin B, Schreiber V, and Dantzer F. Poly(ADP-ribose) polymerases in double-strand break repair: focus on PARP1, PARP2 and PARP3. Exp Cell Res. 2014;329(1):18-25.

26. Murai J, Huang SY, Das BB, Renaud A, Zhang Y, Doroshow JH, et al. Trapping of PARP1 and PARP2 by Clinical PARP Inhibitors. Cancer Res. 2012;72(21):5588–99.

27. Bunting SF, Callen E, Wong N, Chen HT, Polato F, Gunn A, et al. 53BP1 inhibits homologous recombination in Brca1-deficient cells by blocking resection of DNA breaks. Cell. 2010;141(2):243–54.

28. Escribano-Diaz C, Orthwein A, Fradet-Turcotte A, Xing M, Young JT, Tkac J, et al. A cell cycle-dependent regulatory circuit composed of 53BP1-RIF1 and BRCA1-CtIP controls DNA repair pathway choice. Mol Cell. 2013;49(5):872–83.

29. Brambilla E, Moro D, Gazzeri S, and Brambilla C. Alterations of expression of Rb, p16(INK4A) and cyclin D1 in non-small cell lung carcinoma and their clinical significance. J Pathol. 1999;188(4):351-60.

30. Geradts J, Fong KM, Zimmerman PV, Maynard R, and Minna JD. Correlation of abnormal RB, p16ink4a, and p53 expression with 3p loss of heterozygosity, other genetic abnormalities, and clinical features in 103 primary non-small cell lung cancers. Clin Cancer Res. 1999;5(4):791-800.

31. Dean JL, McClendon AK, and Knudsen ES. Modification of the DNA damage response by therapeutic CDK4/6 inhibition. J Biol Chem. 2012;287(34):29075–87.

32. Eleazer R, and Fondufe-Mittendorf YN. The multifaceted role of PARP1 in RNA biogenesis. Wiley Interdiscip Rev RNA. 2021;12(2):e1617.

33. Huang D, and Kraus WL. The expanding universe of PARP1-mediated molecular and therapeutic mechanisms. Mol Cell. 2022;82(12):2315–34.

34. Tempka D, Tokarz P, Chmielewska K, Kluska M, Pietrzak J, Rygielska Z, et al. Downregulation of PARP1 transcription by CDK4/6 inhibitors sensitizes human lung cancer cells to anticancer drug-induced death by impairing OGG1-dependent base excision repair. Redox Biol. 2018;15:316–26.

35. Hopkins TA, Ainsworth WB, Ellis PA, Donawho CK, DiGiammarino EL, Panchal SC, et al. PARP1 Trapping by PARP Inhibitors Drives Cytotoxicity in Both Cancer Cells and Healthy Bone Marrow. Mol Cancer Res. 2019;17(2):409–19.

36. Wei L, Leibowitz BJ, Wang X, Epperly M, Greenberger J, Zhang L, et al. Inhibition of CDK4/6 protects against radiation-induced intestinal injury in mice. J Clin Invest. 2016;126(11):4076–87.

37. Asghar U, Witkiewicz AK, Turner NC, and Knudsen ES. The history and future of targeting cyclin-dependent kinases in cancer therapy. Nat Rev Drug Discov. 2015;14(2):130–46.

