## Supplementary figures and images for "CDK4/6 inhibitors promote PARP1 degradation and act synergistically with PARP inhibitors in non-small cell lung cancer"

### Supplemental figures

**Supplementary Figure 1**

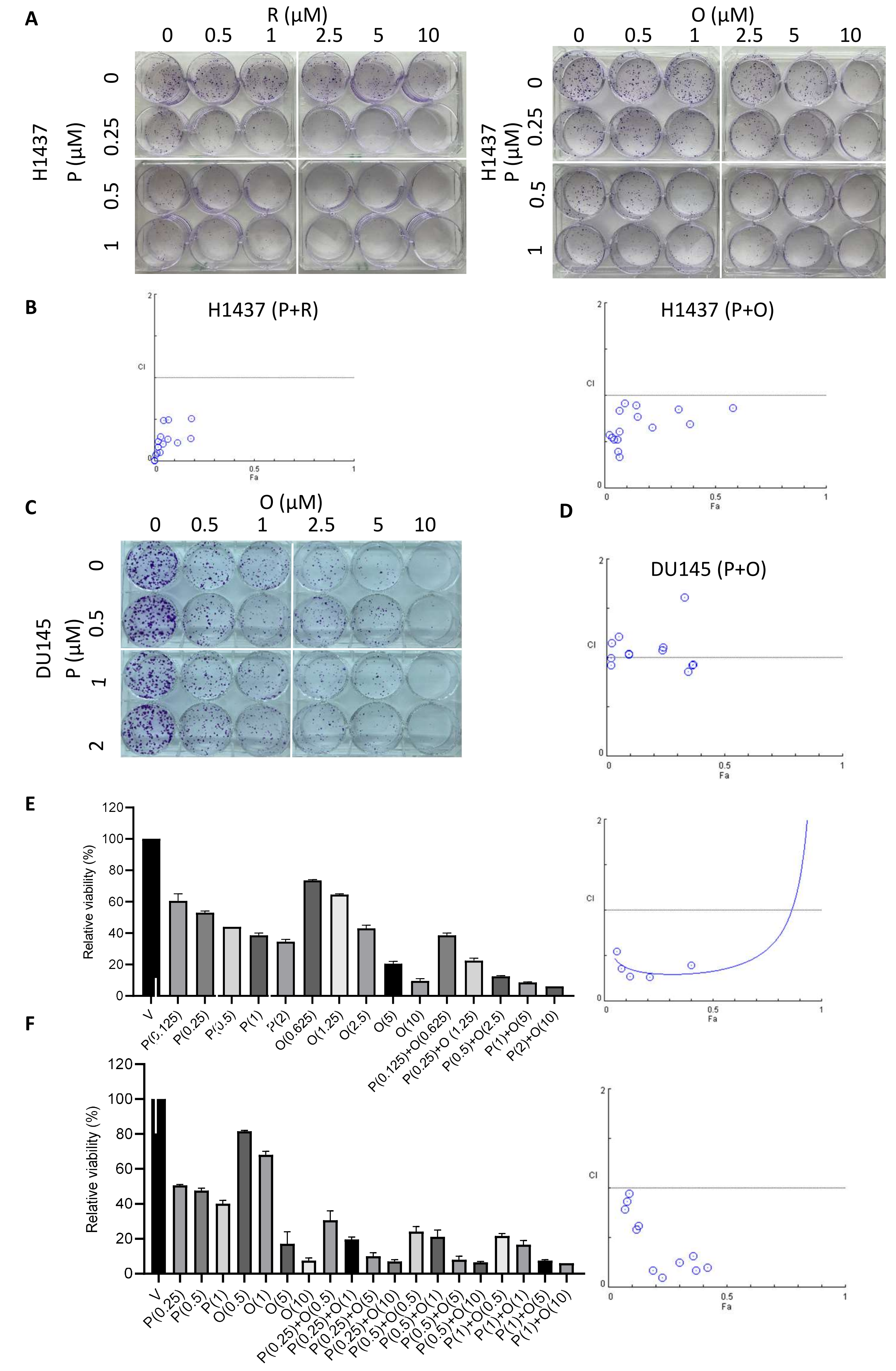

**Supplementary Figure 2**

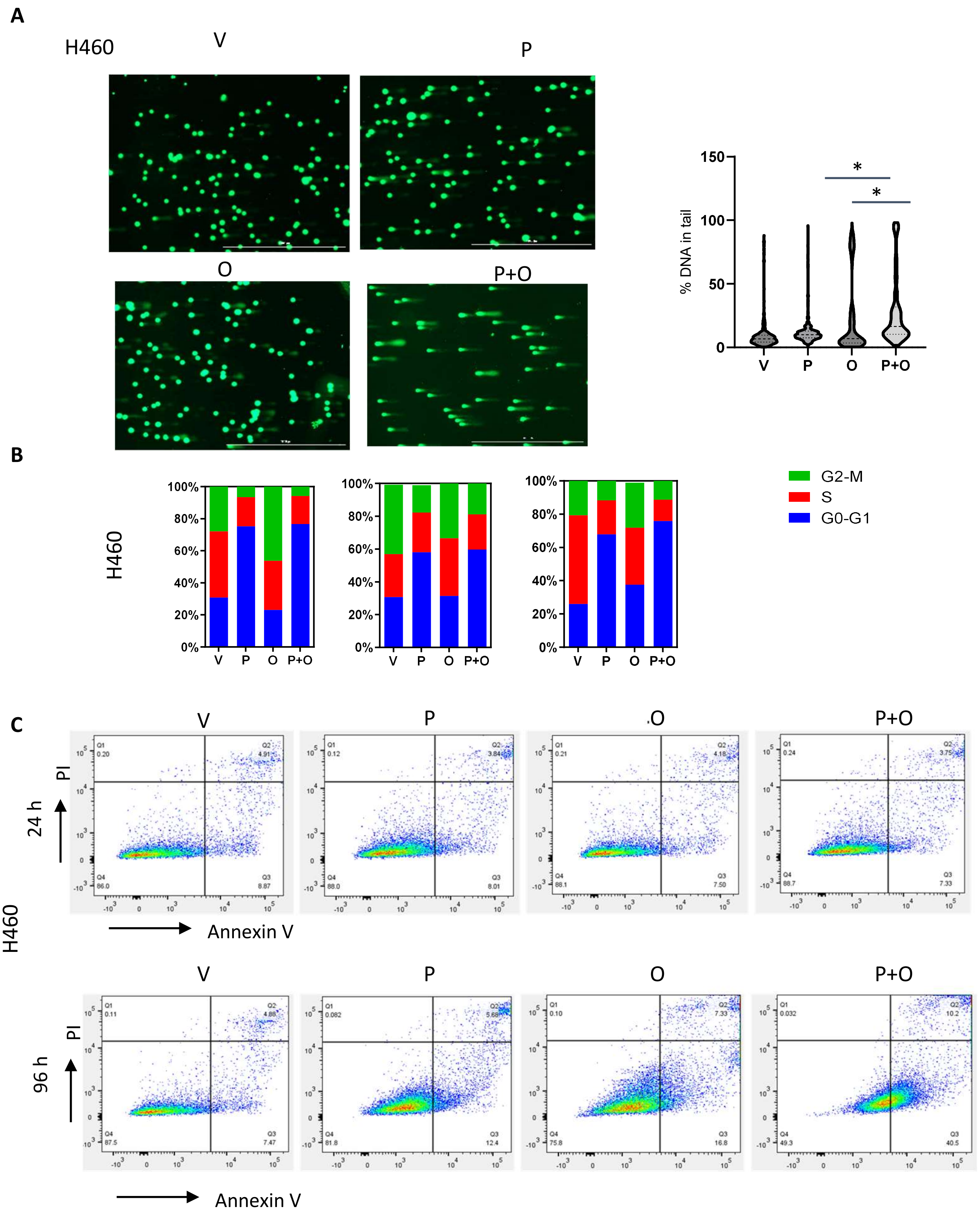

#### Supplementary Figure 3

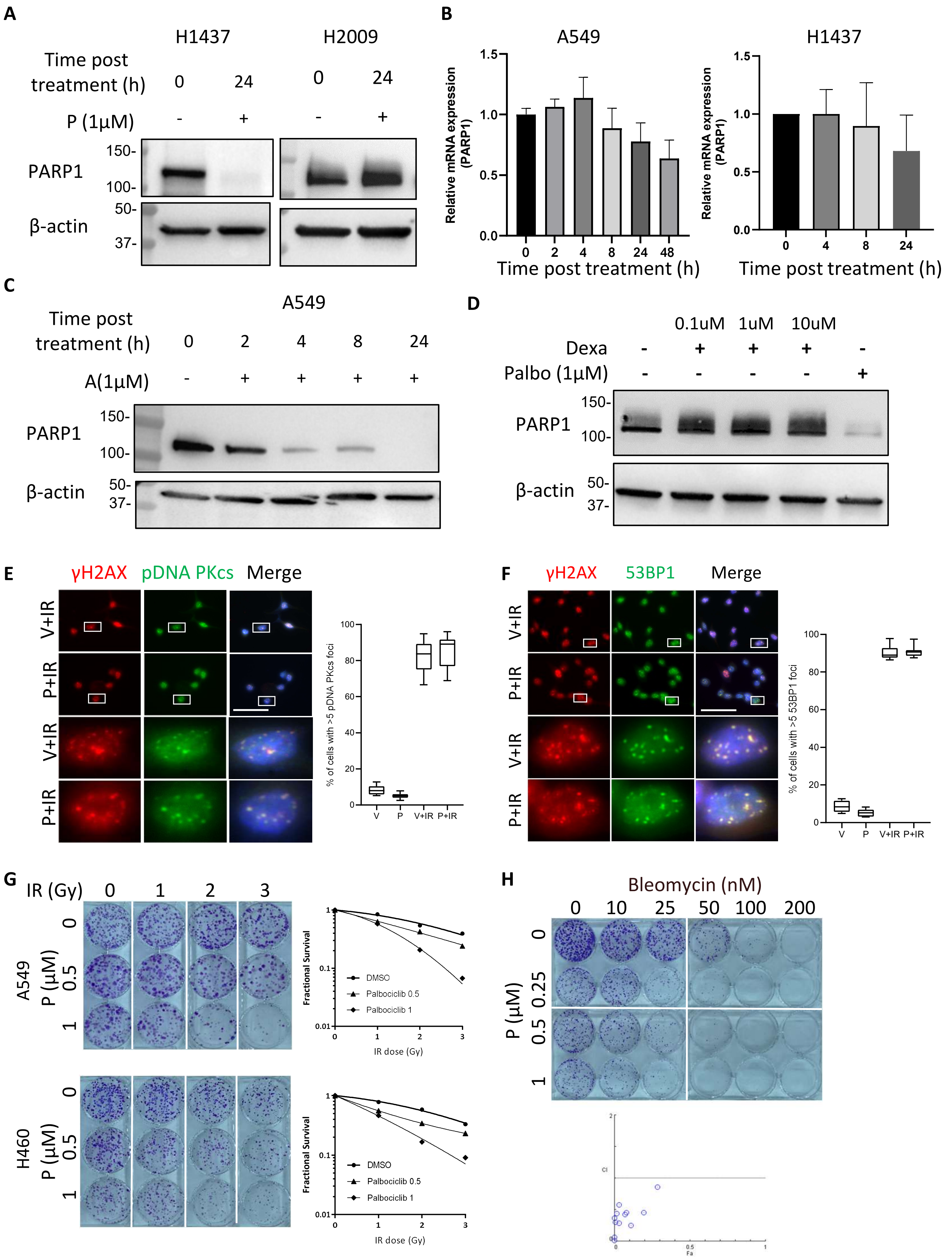

**Supplementary Figure 4.**

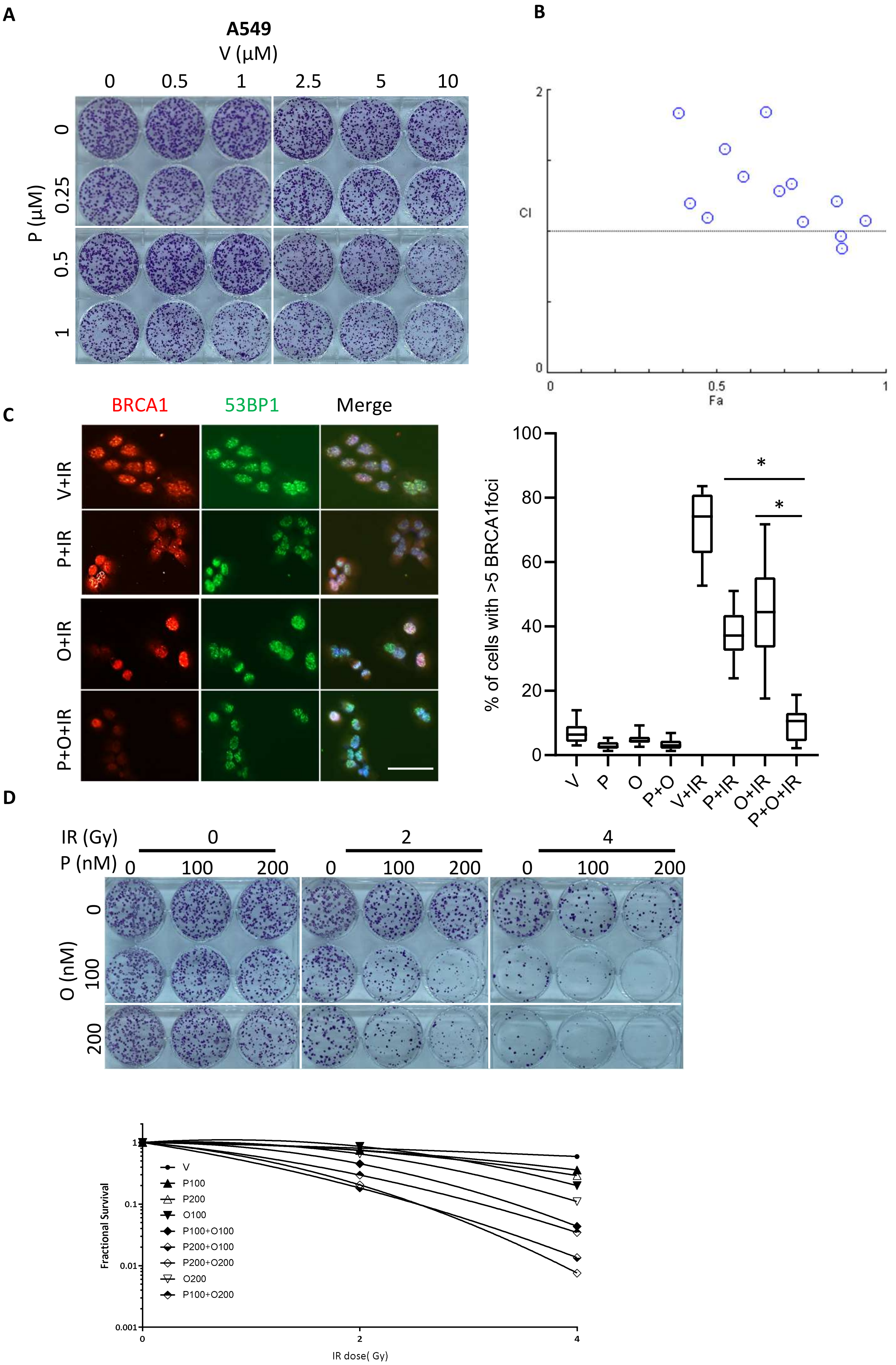
