## Supplemental figure legends for "CDK4/6 inhibitors promote PARP1 degradation and act synergistically with PARP inhibitors in non-small cell lung cancer"

**Supplementary Figure 1. CDK4/6i synergizes with PARPi in RB-proficient NSCLC cell lines but not in RB-deficient cell lines.** (A) Colony formation assays in RB-proficient human NSCLC H1437cell line in the presence of vehicle (V, DMSO), palbociclib (P), olaparib (O)/rucaparib (R), or combination (P+R or P+O). (B) Combination index (CI) values calculated for P+R and P+O from colony formation assay shown in A. All values were less than 1, indicating synergy. (C) Colony formation assays in RB-deficient human prostate cancer cell line DU145 in the presence of vehicle (DMSO), palbociclib (P), olaparib (O), or combination (P+O). (D) CI values of P+O for DU145 cells were mainly more significant than 1, indicating no synergy in the RB-deficient human prostate cancer DU145 cell line. (E) Proliferation assay (left) and CI values (right) in NSCLC A549 cells treated with vehicle (V, DMSO), palbociclib (P, 0.125-2 µM), olaparib (O, 0.625-10 µM), or combination (P+O) at a constant (Dose) ratio (1:5) for 5 days. (F) Proliferation assay (left) and CI values (right) in NSCLC A549 cells treated with vehicle (V, DMSO), palbociclib (P, 0.25-1 µM), olaparib (O, 0.5-10 µM), or combination (P+O) with a non-constant (Dose) ratio for 4 days. Viable cells were quantified by cell viability assays. Data are shown as the mean fold change in cell viability relative to DMSO (mean ± SEM) (n = 3 independent experiments).

**Supplementary Figure 2. CDK4/6i increases PARPi-induced DNA damage in RB-proficient NSCLC cells.** H460 cells were treated with palbociclib (1 μM) and olaparib (2 μM) alone or in combination (P+O) for 72 h. (A) Alkaline comet assay of H460 cells treated with V, P, O, and P+O for 72 h. Representative images of alkaline comet assay (left) and DNA damage quantification by tail moment (right). Box and whisker plots represent values within the interquartile range (boxes) and the minimum to maximum (whiskers). The line within the box shows the median. *P < 0.001 *vs.* single agent treatment, by 1-way ANOVA with Tukey's multiple comparisons test. (B) Flow cytometry analysis of cell cycle distribution in H460 cells treated with V, P, O, or P+O for the indicated time. PARPi increases CDK4/6i induced G1 phase arrest and decreases cell number in the S-phase of the cell cycle. (C) Annexin V/PI double staining apoptosis analysis by flow cytometry in H460 cells treated with V, P, O, or P+O for the indicated time.

**Supplementary Figure 3. CDK4/6i promotes PARP1 protein degradation and regulates DNA repair factor availability and DNA repair competency in RB-proficient NSCLC cells.** (A) Western blot showing that CDK4/6i (palbociclib, P) decreases protein levels of PARP1 in RB-proficient (H1437) but not in RB-deficient NSCLC (H2009) cell line. (B) q-PCR analysis of PARP1 mRNA levels in A549 (left) and H1437 cells (right) treated with palbociclib (P, 1 µM) at indicated time points. CDK4/6i (palbociclib) has no significant effect on mRNA level of PARP1 in A549 and H1437 cells within 24 h treatment. (C) Western blot showing that CDK4/6i (abemaciclib, A) decreases protein levels of PARP1 in a time-dependent manner. (D) Dexamethasone (Dex), a G1 blocker in A549 cells, does not affect protein levels of PARP1 as measured by western blot. (E) CDK4/6i (palbociclib) does not significantly affect IR-induced pDNA PKcs foci recruitment. A549 cells were treated with palbociclib for 16 h and IR (2Gy) for an additional 4 h. The cells were stained with anti-γH2AX and pDNA PKcs antibodies and DAPi and analyzed by immunofluorescence microscopy. (F) CDK4/6i (palbociclib) does not significantly affect IR-induced 53BP1 foci recruitment. A549 cells were treated with palbociclib for 16 h and IR (2Gy) for an additional 4 h, then stained with anti-γH2AX and 53BP1 antibodies and DAPi and analyzed by immunofluorescence microscopy. (G) Colony forming assay showing the radiosensitization effect of CDK4/6i (palbociclib) in RB-proficient A549 and H460 cells. Representative images (left) and quantification (right) were shown. (H) Combination treatment with palbociclib and bleomycin produces synergistic inhibition of clonogenic survival assay in RB-proficient NSCLC A549 cells. Representative images (top) and combination index (CI) values (bottom) were shown.

**Supplementary Figure 4. PARP trapping is essential for the synergy between CDK4/6i and PARPi.** (A) Colony formation assays in RB-proficient NSCLC A549 cell line in the presence of vehicle (DMSO), palbociclib (P), veliparib (V), or combination (P+V). (B) Combination index (CI) values of P+V from the colony formation assay are shown in A. CI values were mainly greater than 1, indicating no synergy for P+V. (C) Impact of the combination treatment of palbociclib and olaparib on IR-induced BRCA1 and 53BP1 foci formation. A549 cells were pretreated with control (DMSO), palbociclib, olaparib, or palbociclib and olaparib combo for 16 h and then exposed to IR (2 Gy) and let them recover for 4 h. Immunostaining was performed using anti-53BP1and anti-BRCA1 antibodies. Representative images (left) and quantification (right) of BRCA1 positive nuclei were shown. Scale bar, 50 μm. (D) Combining a low dose of palbociclib and olaparib sensitizes A549 cells to IR. Colony formation assays were performed on the A549 cells, showing the reduction of cell survival after a low dose of palbociclib/olaparib combined with IR (0, 2, 4 Gy) compared to either treatment alone. Representative images (top) and quantification for fractional survival (bottom) were shown.
